# Metabolic diversity of the Ferrovales and potential contributions to iron oxidation

**DOI:** 10.1101/2025.11.03.686282

**Authors:** Christen L. Grettenberger, Jennifer L. Macalady, Trinity L. Hamilton

## Abstract

Active and abandoned metal and coal mines generate acidic, metal-laden water that pollutes downstream areas, commonly referred to as acid mine drainage (AMD). AMD is host to microbial communities, including acidophilic iron oxidizers. Microbially-mediated iron oxidation is a desirable (bio)remediation strategy for AMD. Ferrovales are a common Fe-oxidizing bacterial group observed in AMD globally and thus could be an exceptional target for bioremediation strategies. However, Ferrovales are difficult to culture and both phylogenetically and metabolically diverse. To better understand the potential for Ferrovales to contribute to AMD bioremediation, we circumvented limitations in culture-based approaches and analyzed 240 genomes and metagenome assembled genomes from the Ferrovales, including taxa from AMD sites with high iron oxidation rates. The phylogenetic and physiological diversity of this group was greater than previously known and multiple Ferrovales taxa can co-occur. For example, we observed taxa that varied in mobility (based on the presence or absence of flagellar biosynthesis genes) and taxa that encode multiple iron oxidation pathways. We also identified Ferrovales that are likely capable of anoxygenic photosynthesis. Our data suggest that differences in physiology promote niche differentiation along resource axes and co-occurrence of multiple taxa. It’s possible that multiple co-occurring iron oxidizing taxa could support a more robust microbial community, resulting in higher iron oxidation rates and more efficient bioremediation.

**Importance:** Acid mine drainage (AMD) pollutes watersheds worldwide. Microbial communities can be leveraged to improve AMD bioremediation, the use of living things for the remediation of contaminated sites, because they drive biogeochemical processes in these ecosystems. Iron-oxidizing microbial species remove iron from AMD effluent, and the iron oxides sorb other trace metals. These communities vary across sites and differ in how rapidly they oxidize iron. Therefore, it is difficult to effectively use them for bioremediation. Ferrovales are common iron-oxidizing taxa in AMD globally, including in sites with exceptionally high rates of iron oxidation. To examine the potential for Ferrovales to be key components of bioremediation strategies, we examined the genomic content and functional potential of Ferrovales in publicly available metagenomic data sets. Our analysis uncovered several new species of Ferrovales as well as an expanded metabolic potential for this group. Comparative genomics suggests that functional diversity may lead to the co-occurrence of multiple Ferrovales species. The presence of multiple iron-oxidizing taxa with distinct physiology could be beneficial for bioremediation strategies.

## Introduction

Acid mine drainage (AMD) is a global pollution problem that pollutes tens of thousands of kilometers of streams worldwide (1–5). AMD is generated when mining activities expose iron-sulfide mineral bearing rocks to oxygen and water. This generates acidic, metal-laden water polluting downstream areas and killing benthic macroinvertebrates, fish, and other aquatic species. Acid mine drainage remediation strategies are costly and often include regular intervention (e.g. active remediation via aeration, lime addition, or bioreactors) (6). Bioremediation is a preferred method for AMD remediation for cost-effective passive processing requiring only minimal continued inputs. One such method leverages the metabolisms of naturally occurring iron-oxidizing species like *Acidithiobacillus ferrooxidans, Gallionella ferruginea*, and *Ferrovum myxofaciens* to oxidize iron, effectively removing it from the stream environment (6). Once iron is removed, the acidity could be neutralized with limestone-lined channels without extensive iron armoring.

The most effective bioremediation systems would oxidize iron rapidly. However, there is significant variation in the iron-oxidation rate at AMD sites. For example, zero-order iron oxidation rates vary by nearly an order of magnitude in Appalachian coal mine drainages and by approximately 5x in the Iberian Pyrite Belt (7). These differences are likely due, in part, to thermodynamics – the Gibbs free energy available from iron oxidation is higher at low pH than it is at higher pH (8). However, thermodynamics cannot fully account for differences in iron oxidation rates. Sites in the Iberian Pyrite Belt had lower iron oxidation rates than those in Appalachian coal mine drainages even when the Gibbs free energy available was greater (7, 8) and chemostats with controlled geochemistry did not replicate field oxidation rate even at similar geochemical setpoints (8, 9). Thus, there is another controller on iron oxidation rates in the field.

To leverage microbial iron-oxidation for bioremediation, we must understand the physiology of iron-oxidizing organisms and the metabolic and ecological factors that correlate with high iron-oxidation rates. Microbial taxa differ in the proteins used to oxidize iron (to transfer electrons from reduced iron (Fe(II)) to the cell) (reviewed in 10) which likely impacts iron oxidation rates. All known iron oxidizers use one of three proteins: Cyc2 like proteins (11, 12), MtoA, or its homolog PioA to oxidize iron (13). Other physiological adaptations may indirectly impact iron oxidation rates by influencing growth and metabolic rates. For example, the ability to fix nitrogen may increase an organism’s growth rate in an environment that is nitrogen limited. Iron oxidizing taxa with flagella may move to ideal redox conditions as the environmental conditions fluctuate across the day/night cycle, resulting in higher net iron oxidation rates compared to immobile taxa. Community-level interactions also likely impact iron oxidation rates. For example, the presence of multiple Fe(II) oxidizing species provides functional redundancy which can stabilize Fe(II) oxidation rates in bioreactors (14) .

The Ferrovales is an iron-oxidizing group commonly found in AMD ecosystems globally (3, 5, 15–26) and contains the genus *Ferrovum*. Ferrovales-dominated communities oxidize iron faster than other communities in field and laboratory-based studies (3, 7, 9). Bioreactors and treatment plants inhabited by *Ferrovum myxofaciens* P3G effectively remove iron from AMD media (25– 28). Therefore, this genus may play a pivotal role in AMD bioremediation worldwide. However, there are multiple clades of Ferrovales in 16S rRNA and genomic data (29, 30). Additionally, published genomes and metagenome assembled genomes have significant genetic diversity. For example, species within *Ferrovum* differ in their ability to fix nitrogen and the presence of genes for flagella (16, 31–34). This likely indicates that different species or strains may have different physiological capabilities and therefore may vary in how they influence geochemistry and thus iron oxidation rates. Furthermore, multiple *Ferrovum* species have been found to co-occur within a single AMD community indicating that strains may take on different roles within an AMD ecosystem (21). Unfortunately, *Ferrovum* is difficult to cultivate and only the type species *Ferrovum myxofacienns* P3G has been isolated (30) limiting culture-based approaches to examine the potential for this group to facilitate AMD bioremediation.

Genomic data can be used to provide insight into the metabolic potential of the Ferrovales. The genome content of *Ferrovum* has been shaped by gene transfer events and genome reduction in some taxa (31) and therefore is not congruent with the evolutionary history of the genus. Thus, it is likely difficult to tie metabolic capabilities to specific taxa using 16S rRNA data alone. Still, to effectively leverage rapid iron oxidation in Ferrovales for AMD bioremediation we must first understand the diversity of physiological traits in Ferrovales taxa and how these traits relate to their phylogeny. To do so, we generated metagenome assembled genomes (MAGs) of Ferrovales spp. from Scalp Level Run (SLR), PA, a site with the highest measured iron oxidation rate in the Appalachian coal belt and Iberian Pyrite Belt (7) with *Ferrovum* spp. as a major component of its microbial community (3). We compared the genome content of SLR *Ferrovum* genomes to nearly 200 publicly available MAGs and genomes of species within the order Ferrovales to investigate the traits of SLR *Ferrovum* spp. that might confer high iron oxidations rates and if these traits are globally distributed.

## Methods

### Scalp Level Metagenomics

A sediment sample was collected from near the emergence of Scalp Level Run, PA (40.25, -78.84). Approximately three grams of surface sediment was collected in a sterile tube, preserved with 15 mL of RNALater, transported on ice, and stored at -20°C. DNA was extracted using a MoBio PowerBiofilm DNA extraction kit (MoBio, Carlsbad, USA). Each sample was extracted twice with a 2- and 4-minute vortex adaptor bead beating time and the products were pooled. Libraries were prepared using the Nextera XT Library Preparation Kit following the manufacturer’s instructions. Paired end 150x150 bp sequencing was performed using an Illumina HiSeq 2500. Reads were quality checked using FastQC 0.11.9 (35). Trimmomatic 0.39 was used to remove leading and trailing low quality and N bases, cut the sequence when the average quality drops below 15, and remove reads shorter than 36 bases long (36). Reads were assembled, using megahit 1.0.6 with a minimum contig length of 1500 bp (37). Reads were mapped using samtools 1.15.1and bowtie2 (38, 39). The jgi-summarize_bam_contig_depths command in metabat 2.12.1 was used to calculate read depth and metabat was used to bin contigs longer than 2500 bp (40). Bins were classified using gtdb-tk 2.1.0 (41). The completeness and contamination for each bin was calculated by CheckM 1.0.13 (42). Bins classified as belonging to the Ferrovales were retained for additional analysis. Metagenomic reads are available on the NCBI Sequence Read Archive under accession number SRX18535481. Metagenome assembled genomes are awaiting NCBI review under submission number SUB15735816. We anticipate that they will be publicly available after the federal workers return from furlough. In the interim, they are available at the Open Science Framework (https://osf.io/zsrw5/overview?view_only=4c3f4d147dd84d999bb7810e98813ec7).

### Pangenomics and Phylogenetics

We searched the NCBI SRA database for all MAGs identified as Ferrovales and retrieved those MAGs (September 25, 2024). Seven genomes identified as *Gallionaceae* spp. were included to provide a root for phylogenetic analyses. Genome completeness and contamination were calculated using CheckM 1.0.13 (42). Because there are disagreements between NCBI and GTDB taxonomies, GTDB-tk 2.1.0 (41) was used to re-classify the MAGs and genomes. Genomes and MAGs that were classified as Ferrovales by both GTDB and NCBI and were more than 75% complete and all MAGs from Scalp Level Run were retained.

Anvi’o v8 was used to generate a contigs database for the Ferrovales bins using the anvi-gen-contigs database command (43). The command anv-run-kegg-kofams annotated putative protein coding genes using the KOfam database (44). The anvi-run-hmms command identified hits the bacteria71 hmm collection, a curated set of single copy genes from GToTree (45) and concatenated and aligned the resulting sequences. IQtree v2.3.1 (46) was used to select the best model for protein substitution and build a consensus tree using 1000 bootstrap replicates for both nonparametric bootstraps and for SH-aLRT. The tree was visualized in the Interactive Tree of Life (iTOL) (47). Genomes were compared using FastANI v1.32 (48) within Anvi’o. Genome similarity was calculated using the anvi-compute-genome-similarity command. Taxa that shared 97% average nucleotide identity (ANI) were retained. FeGenie v1.2 (52) identified iron-related genes in the genomes. Flagellar assembly genes were identified by identifying protein coding genes in the genome using prodigal v2.6.3 (53) and annotating those genes using GhostKoala v3.1 (54) using the prokaryotes + viruses GENES database. Because genomes either contained few (2) or many (>30 out of a potential 54) genes, genomes were considered to encode the genes necessary for flagellar assembly if they contained >30 of the genes in pathway number 20240 (flagellar aseembly). Genes or metabolic pathways were considered to be present if they were present in >50% of the genomes within that species. This approach helps to account for genes that are missing due to the incompleteness of the MAG.

Hidden Markov Models (HMMs) and amino acid sequences for the PufL and PufM reaction center proteins were retrieved from EggNog 5.0 (55). We used these HMMs to identify, retrieve, and translate the genes encoding cyc2, PufL and PufM using the anvi-run-hmms and anvi-get-sequences-for-hmm-hits commands in Anvi’o using a cutoff value of 1E-15. The sequences for PufL and PufM retrieved from EggNog and those from the genomes were aligned using MAFFT v7.505 using the –auto option (56) . The model selected by the software was L-INS-i. The alignment was trimmed with trimAl v1.5.0 (57) using the automated1 method. A phylogenetic tree was built using IQtree v2.3.1. (46) as described above. The trees were visualized in iTOL and rooted at the midpoint for visualization purposes (47).

### Biogeography of Ferrovaceae Species

We used sourmash branchwater (58) to determine the geographic distribution of the phototrophic Ferrovaceae. The representative genome for each species was uploaded to the branchwater metagenome query site hosted by the Joint Genome Institute (https://branchwater.jgi.doe.gov). Branchwater uses sourmash to search 1.2 million publicly available metagenomes and identify those that contained these taxa. To ensure that we were looking only at the hits within the taxon, we removed all hits with ANI less than 97%.

## Results

We retrieved three MAGs from the Scalp Level metagenome. These were 65%-88% complete with 0%-1.2% contamination in 160 – 316 contigs (Supplemental Table 1). We retrieved 240 MAGs identified as Ferrovales from the NCBI SRA. Sixty-one genomes were removed for being less than 75% complete or having more than 5% contamination. One of those genomes was also not identified as Ferrovales by GTDB-tk. The final data set included 179 Ferrovales genomes or MAGs from NCBI, three MAGs from Scalp Level Run, and the seven Gallionaceae genomes used as an outgroup. From these, we identified 22 unique species (≥97% ANI; Supplemental Table 1).

**Table 1.** Metabolic potential of Ferrovales taxa. A single asterisk (*) indicates that the genome for this clade was incomplete (<65%). Therefore, absence in the genome may be due to incompleteness of the MAG. Two asterisks (**) indicate that more than one but fewer than 25% of the genomes in the taxon encode the gene. Empty cells indicate that the taxon does not encode the genes necessary for the metabolic process.

| Clade | Taxon | Genus | Carbon Fixation | Fe(II) oxidation |  | Fe(III) reduction | Nitrogen Fixation | Sulfur oxidation | Anoxygenic Phototrophy | Flagella |
| --- | --- | --- | --- | --- | --- | --- | --- | --- | --- | --- |
|  |  |  |  | Cyc2 | MtoA | MtrB |  |  |  |  |
| 1 | 21 | *Ca. Solisferrum* | yes | yes |  |  |  | yes | yes | yes |
| 1 | 20 | *Ca. Solisferrum* | yes | yes |  |  |  | yes | yes | yes |
| 1 | 19 | *Ca. Solisferrum* | yes | yes | yes | yes |  | yes | yes | yes |
| 2 | 22 | N/A | yes | * |  |  |  |  |  |  |
| 2 | 4 | *Ferrovum* | yes | yes |  |  |  |  |  | yes |
| 2 | 14 | *Ferrovum* | yes | yes | ** |  | yes |  |  | yes |
| 2 | 5 | *Ferrovum* | yes | yes |  |  |  |  |  | yes |
| 2 | 1 | *Ferrovum* | yes | yes |  |  | yes |  |  | yes |
| 2 | 10 | *Ferrovum* | yes | yes |  |  |  |  |  | yes |
| 2 | 8 | *Ferrovum* | yes | yes |  |  |  |  |  | yes |
| 2 | 7 | *Ferrovum* | yes | yes |  |  | yes |  |  | yes |
| 3 | 2 | *Ca. Ferriphilum* | yes | yes |  |  |  |  |  |  |
| 3 | 17 | *Ca. Ferriphilum* | yes | yes |  |  |  |  |  |  |
| 3 | 3 | *Ca. Ferriphilum* | yes | yes |  |  |  |  |  |  |
| 3 | 18 | *Ca. Siderovum* | yes | yes | yes | yes |  |  |  |  |
| 4 | 9 | *Ca. Siderophilum* | yes | yes |  |  |  |  |  | yes |
| 4 | 13 | *Ca. Siderophilum* | yes | yes | yes | yes |  |  |  | yes |
| 4 | 15 | *Ca. Siderophilum* | yes | yes | yes | yes |  |  |  | yes |
| 4 | 11 | *Ca. Acidiferrum* | yes | yes | yes | yes |  |  |  | yes |
| 4 | 12 | *Ca. Acidiferrum* | yes | yes |  |  |  |  |  | yes |
| 4 | 16 | *Ca. Acidiferrum* | yes | yes | yes | yes |  |  |  | yes |
| 4 | 6 | *Ca. Acidiferrum* | yes | yes | yes | yes |  |  |  | yes |

### Diversity of Ferrovales

The MAGs and genomes clustered into four main clades (Figure 1) The first is an outgroup to all other Ferrovales taxa and includes taxa 19, 20, and 21. The second includes eight taxa (taxa 22, 4, 14, 5, 1, 10, 8, and 7) and is an outgroup to the third and fourth clades. The third clade is composed of taxa 2, 17 and 3. The final clade was composed of 8 taxa (18, 9, 13, 15, 11, 12, 16, and 6) respectively. These taxa comprise 7 genera and differ in their metabolic potential. We describe these taxa, propose names for candidate genera, and describe their metabolic potential below. The SLR MAGs are in taxa 22, 1, and 7. The SLR MAG Wind_5 is the only genome in taxon 22. However, it is only 65% complete.

**Figure 1.**
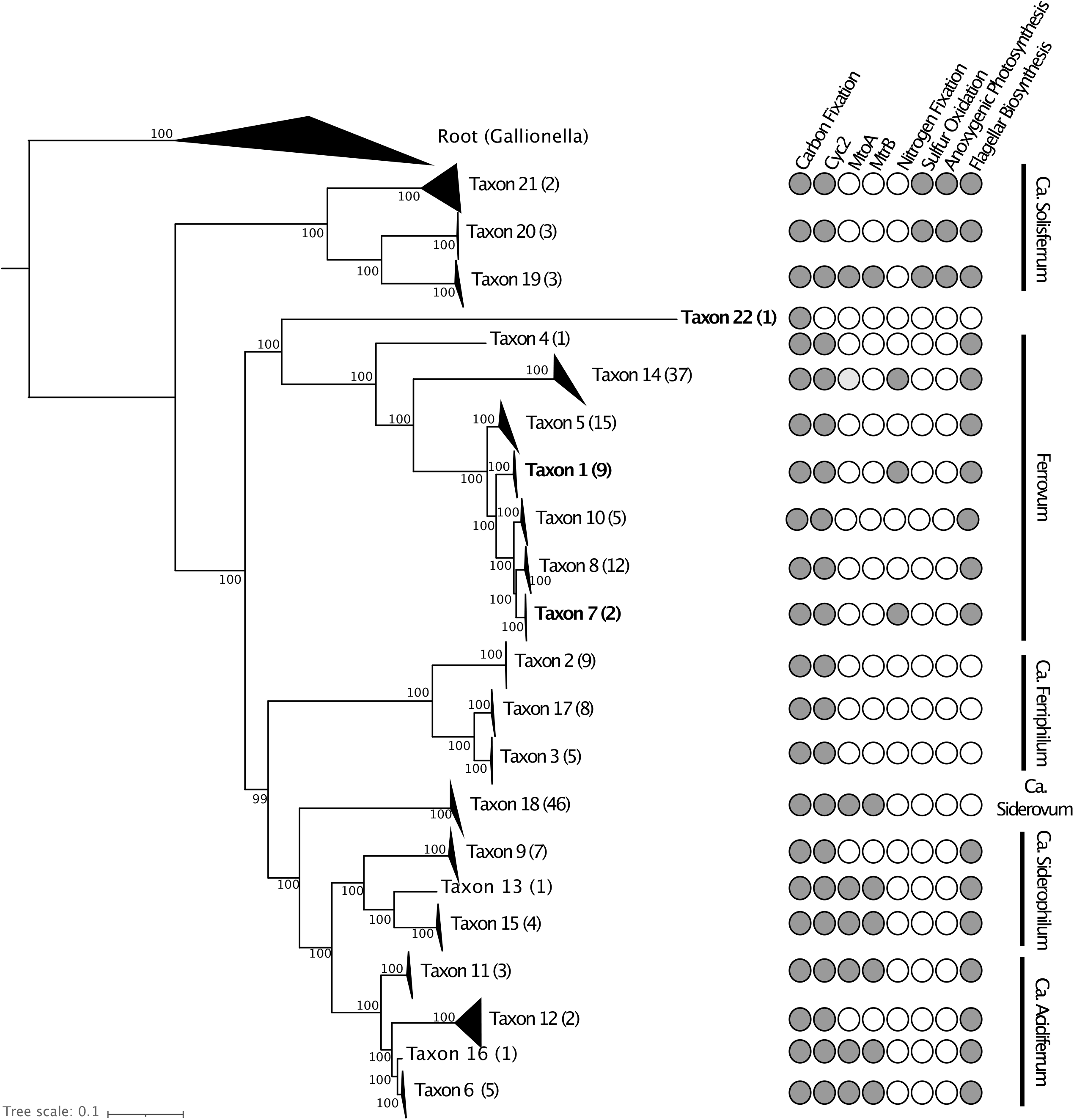
Phylogeny of Ferrovales taxa. Bootstrap values are indicated at the node. The number of genomes present in the taxon is indicated in parentheses. Metabolic potential is indicated by symbols on the leaves.

#### Clade 1

The first clade of Ferrovales is comprised of three taxa from high latitude, stratified lakes (Taxa 19, 20, and 21; Figure 1 No Clade 1 genomes have relatives with more than 97% ANI in any publicly available metagenome outside of the clade. They also share <75% ANI with the other Ferrovales taxa. The ANI cutoff for genera in bacteria is approximately 75% (59), thus Clade 1 taxa are within the family Ferrovaceae but are a separate genus from the other Ferrovales described here. The Genomes Taxonomy Database (GTDB) (60) currently uses the placeholder name CAIVHB01 for this genus. We propose the name “*Candidatus Solisferrum*” gen. nov. (so.lis.fer′rum. L. n. sol, the sun; L. n. ferrum, iron; N.L. neut. n. Solisferrum, referring to the putative ability of this genus to perform photoferrotrophy). All of the members of this clade contain genes encoding the photosystem II reaction center proteins PufL and PufM, carbon fixation via the reductive pentose phosphate cycle, Fe(II) oxidation using Cyc2 and sulfur oxidation from thiosulfate to sulfate. Taxon 19 also encodes the genes necessary for Fe(II) oxidation via MtoA and Fe(III) reduction via MtrB (Table 1). All species encode the genes necessary for flagellar biosynthesis.

#### Clade 2

The second clade of Ferrovales contains 8 taxa (Taxa 22, 4, 14, 5, 1, 10, 8, and 7) from AMD sites across the United States, Europe, and China. This clade contains two genera including the genus *Ferrovum* (Taxa 4, 14, 5, 1, 10, and 7) and an additional unnamed taxon from SLR (Taxon 22). The SLR MAG is only 65% complete which might confound accurate phylogenetic placement. Members of this clade are capable of carbon fixation via the reductive pentose phosphate cycle and Fe(II) oxidation with Cyc2. Three (taxa 1, 7, and 14) are capable of nitrogen fixation (Table 1). All species except Taxon 22 encode the genes necessary for flagellar biosynthesis.

#### Clade 3

The third clade of Ferrovales contains three taxa (Taxa 2, 17, and 3) from mining waste and AMD sites across Canada, Europe, and China. This clade includes the named species *Ferrovum* sp. PN-J185. Based on an ANI cutoff of 75% and GTDB-tk results, this clade is a separate genus from the other Ferrovales discussed here. Therefore, we propose the name “*Candidatus Ferriphilum*” gen. nov. (fer.ri.’phi.lum. L. n. ferrum, iron; Gr. Adj. philos, loving; N.L. neut. n. Ferriphilum, referring to the preference of this genus for iron-containing environments and putative ability to perform iron oxidation). This name would replace the GTDB placeholder genus name PN-J185. Members of this clade are capable of carbon fixation via the reductive pentose phosphate cycle and Fe(II) oxidation with Cyc2. One taxon (Taxon 18) is capable of Fe(II) oxidization using MtoA and Fe(III) reduction using MtrB (Table 1). None of the species encode the genes necessary for flagellar biosynthesis.

#### Clade 4

The final clade of Ferrovales contains eight taxa (Taxa 18, 9, 13, 15, 11, 12, 16, and 6) and includes taxa from stratified lakes in Finland (Taxon 18), and AMD sites in China, the United States. Based on an ANI cutoff of 75% and GTDB-tk results, this clade contains three genera currently indicated by GTDB placeholder names CAIVUX0 (Taxon 18), JAKBAT01 (Taxa 9, 13, and 15) and JAJZRV01 (Taxa 16, 6, 11 and 12).

To replace placeholder name CAIVUX0 (Taxon 18), we propose the name “*Candidatus Siderovum* gen. nov, (sid.er.o’vum Gr. N. sideros, iron; L.n. ovum, egg; N.L. neutr. N. Sideovum, “iron egg” referring to the preference of this taxon for iron-rich environments and referencing the cell shape described in the *Ferrovum* type species).

To replace placeholder name JAKBAT01 (Taxa 9, 13, and 15), we propose the name “*Candidatus Siderophilum*” gen. nov. (sid.er.o’.phil.um Gr.n sideros, iron; Gr. Adj. philos, loving; N.L. neut. n. Siderophilum, referring to the preference of this genus for iron containing environments and putative ability to perform iron oxidation).

To replace placeholder name JAJZRV01 (Taxa 16, 6, 11 and 12), we propose the name “*Candidatus Acidiferrum* gen. nov.(acidi.ferr’um ; L.adj. acidus, acidic; L.n. ferrum, iron; referring to the preference of this organism for acidic environments and its putative ability to perform iron oxidation).

Members of this group are capable of carbon fixation via the reductive pentose phosphate cycle, Fe(II) oxidation with Cyc2. Five taxa (Taxa 13, 15, 11, 16, and 6) are capable of Fe(II) oxidation with MtoA and Fe(III) reduction with MtrB (Table 1). All species except Taxon 18 encode the genes necessary for flagellar biosynthesis.

### Anoxygenic Photosynthesis

Three of the taxa encode genes for anoxygenic photosynthesis via photosystem II including the PufL and PufM subunits and are therefore likely capable of anoxygenic photosynthesis. All genomes from this taxon also encode the PufL and PufM subunits. The translated sequences from these taxa form a monophyletic group in the phylogenies of pufL and pufM (Figure 2) with strong bootstrap support of their node (bootstrap support = 99 for PufL and 100 for PufM). The closest relatives to the PufL sequences are a clade of sequences from the alphaproteobacterial; genera *Rhodopseudomonas, Bradyrhizobium, Hypohomicrobium, Mesorhizobium*, and *Rhodospirillum* though this node has weaker bootstrap support (bootstrap = 59) (Figure 2A) The closest relatives to the PufM sequences are a clade of sequences from the alphaprotobacterial genera *Magnetospirillum, Acidiphilium, Hoeflea, Rhoduvulum, Salipiger, Roseivivax, Roseobacter, Oceanicola, Roseovarius, Paracoccaceae, Roseicyclus, Jannaschia, Dinoroseobacter*, and *Thalassobacter*, the betaproteobacterial order *Limnohabitans*, and gammaproteobacterial genus *Ectothiorhodospira* (Figure 2B). The overall phylogenetic placement of these groups was retained in a tree that was constructed with both PufL and PufM encoding genes (data not shown).

**Figure 2.**
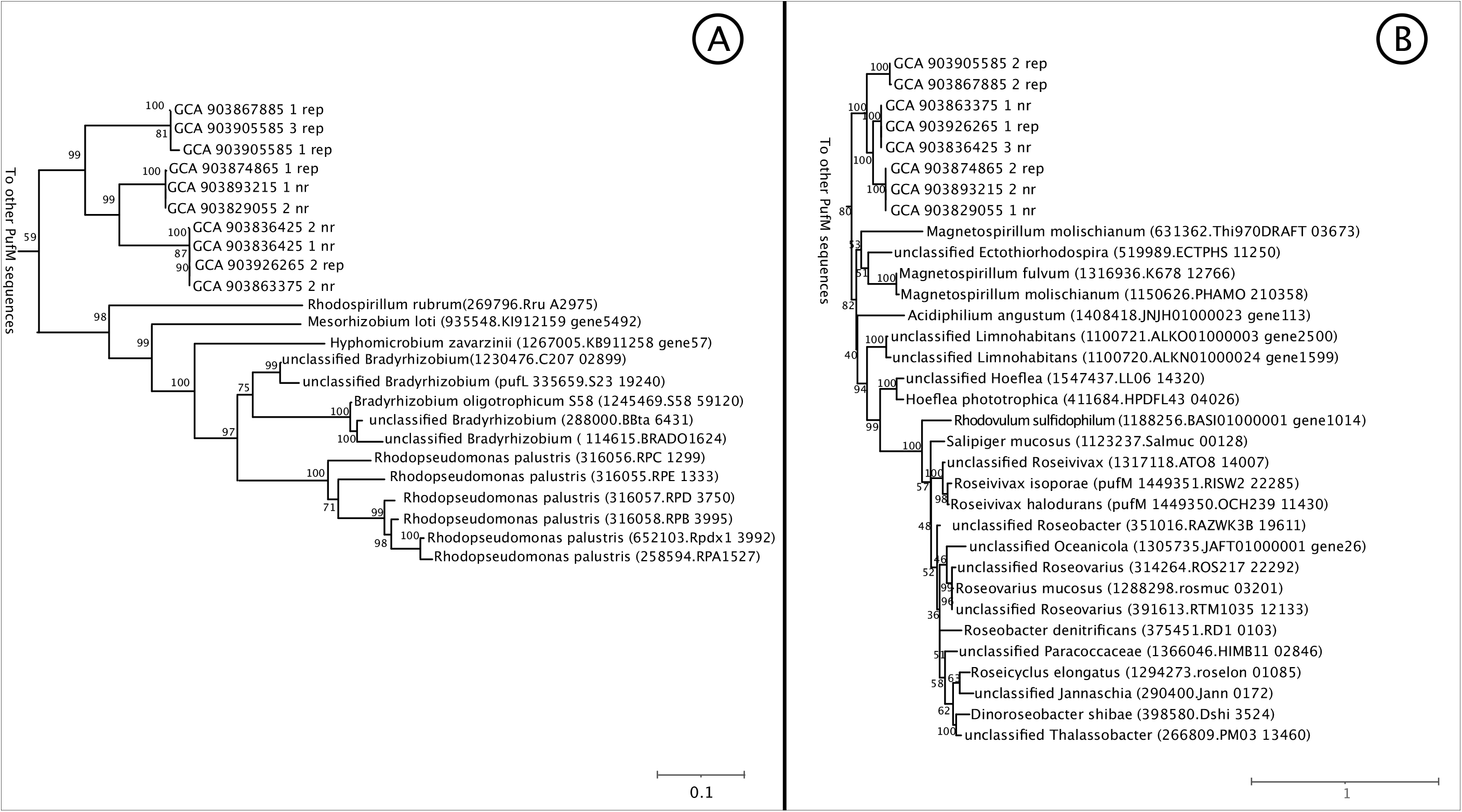
Phylogeny of A) PufL and B) PufM translated nucleotide sequences. Trees were rooted at the midpoint for visualization purposes. Sequences from each taxon are indicated in bold with the accession number for the genome in parentheses.

## Discussion

### Diversity of the Ferrovales

The diversity of the Ferrovales is higher than previously reported (e.g., 16, 21, 31) both phylogenetically and metabolically Our analysis shows that the Ferrovales comprise four clades with 7 genera and 22 species. These taxa share some metabolic traits; all species appear to be capable of iron oxidation using the Cyc-2 like protein and carbon fixation via the reductive pentose phosphate. However, they differ in their ability to fix nitrogen, use alternative Fe(II) oxidation pathways, reduce iron, oxidize thiosulfate, and perform anoxygenic phototrophy.

#### Anoxygenic Photosynthesis

Three of the taxa (taxa 19, 20, and 21) encode genes for the PufL and PufM photosystem II proteins. To our knowledge, the capacity for anoxygenic phototrophy has never been reported in the Ferrovales before but is common in the Proteobacteria including the Burkholderiales, close relatives of the Ferrovales. Because these species are capable of iron oxidation, it is likely that they perform photoferrotropy or the use of Fe(II) as an electron donor in anoxygenic photosynthesis. Photoferrotrophy is reported in a wide variety of groups including the Proteobacteria (e.g. *Rhodovulm*, Rhodopseudomonas, and Rhodobacter spp.), Chlorobi (e.g. *Chlorobium* spp.) (e.g., 61–63). Alternately, they use thiosulfate as an electron donor for anoxygenic photostophy similar to the green and purple sulfur bacteria (Chlorobi and Chromatiaceae and Ectothiorhodospiraceae) (64). Alternately, these species encode the genes sulfur oxidation and therefore may using thiosulfate as an electron donor. The reaction center sequences from the Ferrovales are monophyletic and closely related to sequences from alpha and gamma proteobacteria suggesting that their shared ancestor acquired the genes for these proteins via a single horizontal transfer event.

The three taxa capable of anoxygenic phototrophy inhabit stratified freshwater lakes in high latitudes (65) . Based on samples with available site descriptions, these taxa reside at depths that putatively in the photic zone (0 – 0.7 meters). At the 97% similarity level, Branchwater did not identify any additional environments in which these taxa were found including AMD. Because Branchwater finds all instances of these taxa in publicly available datasets (>1 million metagenomes), this suggests that they may be geographically restricted, or dispersal limited.

#### Iron Cycling

Iron oxidizing taxa use iron as an electron source while also assimilating iron for cellular processes. For electrons, iron oxidizing taxa use outer membrane proteins to oxidize Fe(II). All known acidophilic iron oxidizers oxidize iron using cytochrome *c* in the outer membrane and the electrons are passed on through the periplasm and inner membrane using a variety of redox proteins (reviewed in 10). Neutrophilic iron oxidizers have a wider variety of iron oxidizing complexes including MtoA and its homolog PioA which also oxidize iron in the outer membrane (13). Based on our analyses, Ferrovales differ in strategies for iron oxidation. All recovered Ferrovales MAGs encode the genes for Cyc2 iron oxidation while a subset also encodes the genes for the MtoA pathway. Although both the Mto and Cyc2 complexes oxidize iron, they may be used under different geochemical or environmental conditions. The Cyc2 complex is smaller than Mto and is predicted to be a single heme (12). Therefore, synthesis and maintenance of the Cyc2 complex may require fewer resources (12). However, Mto can oxidize mineral-bound Fe(II) (66) in addition to dissolved Fe(II) suggesting that the taxa that encode Mto may be able to use solid Fe(II). Oxidizing different forms of Fe(II) could allow Mto-encoding taxa to persist in microenvironments where dissolved Fe(II) concentrations are low, potentially due either to limited source or high competition (or both). The MtoA complex is typically used by neutraphilic iron oxidizers where Fe(II) concentrations are generally lower. Therefore, the taxa that encode MtoA may have a competitive advantage in low iron or more neutral pH environments. Several taxa also encode the genes necessary for MtrB. MtrB was found in dissimilatory iron reducing taxa including *Shewanella*, and in these species it is used to transport electrons across the membrane to solid Fe(III). If MtrB in Ferrovales is used for dissimilatory iron reduction, it would provide a competitive advantage in anaerobic environments where Fe(II) is unavailable, but poorly crystalline Fe(III) minerals are abundant like in drying sediments. Alternatively, it may be used in microenvironments with reducing conditions.

### Contribution of Ferrovales to Rapid Iron Oxidation at Scalp Level Run

The rapid iron oxidization rate at SLR may be driven by a number of biotic and abiotic factors. Fe(II) oxidation is likely determined in part by the composition of the microbial community as well as pH and oxygen and Fe(II) concentration (19, 67). At SLR, pH is low (2.77-2.96) (3) and Fe(II) oxidation is more energetically favorable at low pH (8). However, Fe(II) oxidation at SLR is higher than at other sites with similar pH (8) indicating rates are not controlled by pH alone. Compared to sites with similar physiochemical conditions, the microbial community at SLR is distinct. At SLR, *Ferrovum* accounts for 49 – 63% of the microbial community suggesting that these Fe(II) oxidizing taxa contribute to rapid Fe(II) oxidation rates (3).

Increased diversity increases the number of ecosystem functions and services provided by a microbial community in terrestrial ecosystems (68). Thus, the genetic and inferred physiological diversity provided by multiple species of *Ferrovum* could contribute to the rapid rate of Fe(II) oxidation at Scalp Level Run. Three *Ferrovum* taxa inhabit the single location sampled at Scalp Level Run. All three taxa found in SLR are members the second clade (Figure 1) and appear to have largely similar metabolic potential. Two taxa, within Taxa 1 and 7, encode the genes necessary for nitrogen fixation and flagellar biosynthesis while the MAG in Taxon 22 does not, but this may be due to genome incompleteness. These taxa may provide functional redundancy at Scalp Level run or they may partition the Fe(II) oxidizing niches in microenvironments at SLR and other locations where multiple species of Ferrovale*s* co-occur.

Bioreactors inoculated by sediment from Scalp Level Run oxidized Fe(II) at rates lower than in situ (at SLR) (9) . However, only two (Taxa 7 and 5) were detected in the bioreactors (14, 69). The increased diversity of Ferrovales in SLR may enable the iron oxidizing community to oxidize iron in different niches, under different geochemical microenvironments, or at different times of the day (e.g. as the concentration of oxygen varies based on the activity of eukaryotic phototrophs). This could have the net effect of increasing iron oxidation rates by maximizing the locations where and periods of time during which that iron can be oxidized, consistent with higher rates of Fe(II) oxidation compared to similar sites. Co-occurring species of *Ferrovum* have been observed in other AMD sites (e.g. Cabin Branch in Kentucky) (21) but Fe(II) oxidation rates have not been reported. Unfortunately, very few sites have both a measured iron oxidation rate and microbial community composition, so this hypothesis (that co-occurring *Ferrovum* result in higher Fe(II) oxidation rates) cannot be tested with currently available data. Alternately, some taxa of Ferrovales may oxidize Fe(II) more rapidly than others, and their presence may increase Fe(II) oxidation rate as a whole. However, testing this hypothesis would require culture-based experiments and unfortunately *Ferrovum* species are extremely difficult to culture (30).

The diversity of the entire guild of iron oxidizers, rather than just within the Ferrovales, helps to determine the iron oxidation rate. As with Ferrovales taxa, multiple co-occurring Fe(II) oxidizers may inhabit distinct niches that are not distinguishable using modern field methods. These co-occurring species may allow Fe(II) oxidation to proceed in more microenvironments and provide stability when geochemical conditions change (14). At Scalp Level Run, members of Ferrovales predominate, making up >50% of the microbial community (3). However, *Ferrovum* is not the only iron oxidizer present. 16S rRNA data suggest the presence of other iron-oxidizing taxa including an AMD specific clade of Xanthomonadales most closely related to the cultivated taxon WJ2 (3) and the physiological diversity of these Fe(II) oxidizers may contribute to the rapid Fe(II) oxidation rate. Multiple iron-oxidizing taxa regularly co-occur in AMD suggesting both intra- and interspecific diversity are key factors for microbially-mediated iron oxidation. Effective bioremediation strategies should consider the potential for taxonomic and physiological diversity to contribute to more efficient bioremediation. Future studies could best inform bioremediation strategies by combining analysis of microbial community composition and function along with subsequent in situ Fe(II) oxidation rates.

## Acknowledgements

This work was funded in part through US National Science Foundation grant No. 2345568 to Grettenberger and Hamilton. Field work and sequencing were funded by the Office of Surface Mining and Reclamation under grant number S11AC20005 to Macalady and a Geological Society of America Graduate Student Research Grant to Grettenberger.

---

**Supplemental Table 1**. Genomes used in the pangenome analysis, taxonomic classification, completeness, contamination, and strain heterogeneity of each genome.

